# Mesenchymal Stem Cell-Derived Extracellular Vesicles Mitigate Immune Cell Activation in an In Vitro Model of Post-Resuscitation Inflammation

**DOI:** 10.1101/2025.02.13.637856

**Authors:** Tyler J Rolland, Sumbule Zahra, Daniel Cucinotta, Rebeccah Young, Brian Weil

## Abstract

**Background:** Systemic inflammation is a well-established component of post-cardiac arrest syndrome (PCAS), a condition responsible for significant morbidity and mortality in patients who are initially resuscitated from sudden cardiac arrest. Mesenchymal stem cell-derived extracellular vesicles (MSC-EVs) have emerged as promising immunomodulatory agents in various inflammatory conditions, including after ischemia-reperfusion injury (IRI). Here, we investigated the therapeutic potential of MSC-EVs in porcine peripheral blood mononuclear cells (PBMCs) stimulated with lipopolysaccharide (LPS) or mitochondrial DNA (mtDNA) to mimic immune cell activation in PCAS.

**Methods:** PBMCs were isolated from healthy pigs (*Sus scrofa*), cultured *in vitro*, stimulated with LPS or mtDNA, and treated with a range of MSC-EV concentrations. Flow cytometry, quantitative PCR, ELISA, and ROS/RNS measurements were performed to assess PBMC activation.

**Results:** MSC-EV treatment reduced LPS-induced inflammatory granulocyte activation and selectively modulated cytokine transcripts, including IFNα, IL-1β, and TNF-α, in a concentration-dependent manner. Similar immunosuppressive effects were observed in mtDNA-stimulated PBMCs, where MSC-EVs attenuated dendritic cell activation and inflammatory cytokine release. Furthermore, higher concentrations of MSC-EVs significantly decreased ROS/RNS production in both LPS- and mtDNA-challenged PBMCs.

**Conclusions:** MSC-EVs exhibit potent immunomodulatory properties against LPS- and mtDNA-induced activation of porcine PBMCs, highlighting their broad capacity to modulate immune responses and mitigate oxidative stress induced by pro-inflammatory stimuli that are relevant to PCAS. These findings provide further support for the administration of MSCs, or MSC-EVs themselves, as a potential therapeutic intervention to target immune activation in PCAS and other disorders characterized by an acute systemic inflammatory state.

## INTRODUCTION

Systemic inflammation is a critical contributor to morbidity and mortality in post-cardiac arrest syndrome (PCAS), a condition that arises after successful return of spontaneous circulation (ROSC) from sudden cardiac arrest (SCA) (1, 2). Within the first 24-48 hours post-ROSC, patients commonly exhibit “sterile sepsis,” characterized by an increase in circulating concentrations of lipopolysaccharide (LPS) and marked immune cell activation (3). In addition to LPS, recent evidence from our laboratory and others (4-8) implicates mitochondrial DNA (mtDNA) as a potent pro-inflammatory stimulus in PCAS and other conditions characterized by ischemia-reperfusion injury (IRI). Because LPS and mtDNA each contribute to the heightened inflammatory state observed after resuscitation from SCA, novel interventions that modulate immune cell activation by these stimuli may offer therapeutic benefit in patients with PCAS (9).

Mesenchymal stem cells (MSCs) have gained attention as potential immunomodulatory agents in a variety of inflammatory conditions, including preclinical models of IRI (e.g., stroke (10), acute myocardial infarction (11), acute kidney injury (12), and porcine models of PCAS (13)). Although MSC-based therapies have shown promise, their efficacy remains modest, with accumulating evidence suggesting that MSC-derived extracellular vesicles (MSC-EVs) may account for much of MSCs’ beneficial effects (14-18). Previous investigations highlight MSC-EVs’ cytoprotective and immunomodulatory properties; for example, EVs derived from MSCs can diminish infarct size (19), enhance neovascularization (20), reduce fibrosis (21, 22), and improve cardiac function after myocardial IRI (23-27). These beneficial effects are generally attributed to the paracrine capacity of MSC-EVs, which can deliver bioactive molecules capable of modulating immune activation and cytokine release.

Despite this emerging paradigm, the precise mechanisms by which MSC-EVs regulate immune cells, particularly in the context of activation by potent stimuli like lipopolysaccharide (LPS) and mitochondrial-DNA (mtDNA), remain incompletely understood. Accordingly, we investigated the therapeutic potential of MSC-EVs to mitigate inflammatory responses in naïve porcine peripheral blood mononuclear cells (PBMCs) stimulated with LPS or mtDNA to mimic immune cell activation in PCAS. Using multiple concentrations of MSC-EVs to determine if putative effects were dose-dependent, we evaluated pro-inflammatory phenotypic shifts (via flow cytometry), transcript expression (qPCR), cytokine secretion (ELISA), and ROS/RNS production in response to MSC-EV treatment. The results provide novel insight regarding the ability of MSC-EVs to attenuate immune cell activation induced by LPS and mtDNA, thereby supporting further investigation of MSC-EVs as a therapeutic intervention for PCAS and other conditions characterized by an acute systemic inflammatory response.

## METHODS

### Ethics Statement

All protocols followed institutional guidelines for animal research and were approved by the State University of New York at Buffalo Institutional Animal Care and Use Committee. Investigators were blinded to conditions (e.g., mtDNA vs. mtDNA + MSC-EVs) while conducting analytical protocols and analyzing data. Please see **Supplemental Materials** for complete methodological details.

### MSC Isolation and Culture

Bone marrow-derived mesenchymal stem cells (MSCs) were obtained from bone marrow of healthy White Yorkshire × Landrace swine and cultured in Advanced DMEM with Glutamax, 10% FBS, and penicillin/streptomycin. At passage zero, cells were cryopreserved in a supplemented medium containing B-27 Supplement and DMSO. The full protocol is provided in the **Supplemental Materials**.

### MSC-EV Isolation and Characterization

Revived MSCs were maintained in serum-free medium for three days, and the conditioned medium was collected. Extracellular vesicles (EVs) were isolated by sequential low-speed centrifugation, 0.2 μm filtration, and concentration using 30 kDa cutoff centrifugal filters. Particles were characterized by ZetaView® particle tracking analysis and verified with the Exo-Check™ Exosome Antibody Array. Details are available in the **Supplemental Materials**.

### PBMC Isolation and Culture

Peripheral blood mononuclear cells (PBMCs) were isolated from whole blood of healthy pigs using Ficoll separation and RBC lysis. Cells were cultured in RPMI 1640 supplemented with 10% FBS and antibiotics. After 1 week, PBMCs were activated for 24 hours with 1× LPS or 1 μg/mL mtDNA in lipofectamine-based transfection reagent, with MSC-EVs added in place of FBS. (**Figure S1**; see **Supplemental Materials** for additional information).

### Flow Cytometry

Cultured PBMCs were labeled with porcine-specific surface markers for inflammatory phenotyping and stained with 7AAD for viability (**Figure S2)**. Data were acquired on a BD LSRFortessa™ and analyzed using FCS Express 7 Plus. MSC-EV uptake was assessed via Vybrant DiD labeling. (**Table S1**; see **Supplemental Materials** for additional information).

### qPCR

Total RNA was isolated (RNeasy Kit) and quantified by NanoDrop. cDNA synthesis was followed by qPCR with primers for IFNα, IL-1α, IL-1β, IL-6, IL-8, TNFα, and β2M, using a SYBR Green Supermix. Expression levels were normalized to β2M. (**Table S2**; see **Supplemental Materials** for additional information).

### ELISAs and ROS/RNS Assays

Conditioned media (CM) were clarified by centrifugation before being assessed by porcine-specific ELISAs against inflammatory cytokines, while ROS/RNS production was evaluated using a DCF-based kit. (**Table S3**; see **Supplemental Materials** for additional information).

### Statistical Analysis

Data are presented as mean ± SEM. Two-tailed unpaired Student’s t-tests were used to compare groups; concentration-response was assessed by simple linear regression of responses to multiple concentrations of MSC-EVs. Significance was set at p<0.05. Analyses were conducted with GraphPad Prism Software. Additional details are included in the **Supplemental Materials**.

## RESULTS

### MSC-EV Characterization

ZetaView analysis of isolated MSC-EVs indicated a predominantly spherical population of vesicles with a mean diameter of approximately 100 nm (**Figure 1A and B**). This size distribution and morphology align with standard EV characterizations, reinforcing the reliability of our isolation technique (28). Further validation using the Exo-Check exosome antibody array revealed robust expression of established EV markers, including CD63, EpCAM, ANXA5, TSG101, GM130, FLOT1, ICAM, ALIX, and CD81. These markers were readily detected in both 100μg and 1μg sample groups, confirming the presence of EVs (**Figure 1C and D**). Collectively, these findings demonstrate the successful isolation and characterization of MSC-EVs, providing a rigorous foundation for subsequent functional assays.

**Figure 1.**
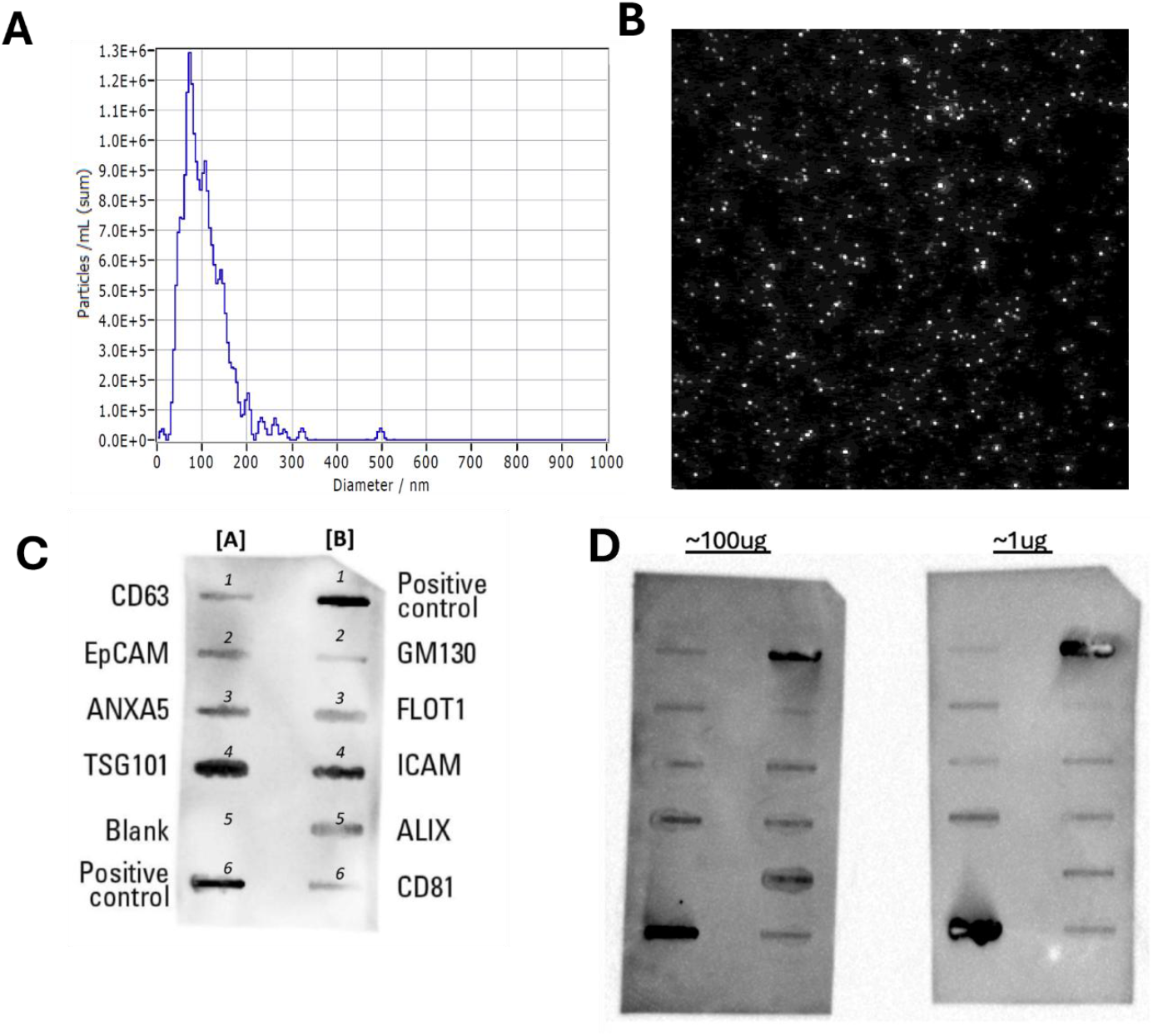
Porcine MSC-EV Validation and Characterization. A) Representative histogram depicting the diameter distribution of MSC-EVs from conditioned media following ZetaView analysis indicate particle size of ∼100 nanometers. B) Representative ZetaView image illustrating the circular morphology of the MSC-EVs. C) ExoCheck antibody layout used as a comparison for figure 1D. D) ExoCheck blots show the presence of surface markers in line with extracellular vesicles in MSC-EV samples.

### Modulation of LPS-Triggered Inflammatory Cell Phenotype by MSC-EVs

To evaluate the capacity of MSC-EVs to mitigate LPS-induced immune activation, flow cytometric analyses were performed on LPS-stimulated PBMCs incubated with DiD-labeled MSC-EVs at various concentrations. MSC-EV treatment (2e9 EVs) exhibited a trend towards a reduction of LPS-driven inflammatory phenotype shift in CD172+ cells (**Figure 2A and S3A**), primarily due to a decrease in the CD172+CD163− inflammatory granulocyte subpopulation (**Figure 2C and S3C**) that was driven by altered surface marker expression of cells that did not take up MSC-EVs. In contrast, the proportions of CD172+CD16+ dendritic cells (**Figure 2B and S3B**) and CD14+CD163+ macrophages (**Figure 2D and S3D**) remained largely unchanged compared with LPS-only controls which did not elicit strong inflammatory activation of these populations at 24hrs post-activation. Furthermore, albeit not significant, both MSC-EV-uptake and no-EV-uptake groups exhibited comparable immunophenotypic profiles within granulocytes in a concentration-dependent manner (**Fig S3 and Table S4**). Notably, these effects occurred in the absence of cell death (**Figure S4A**). Apart from the observed reduction in inflammatory granulocytes, MSC-EVs did not significantly modulate other immune cell subsets. This suggests that MSC-EVs selectively modulate LPS-induced inflammatory granulocyte activation without affecting broader immune cell populations under these *in vitro* conditions.

**Figure 2:**
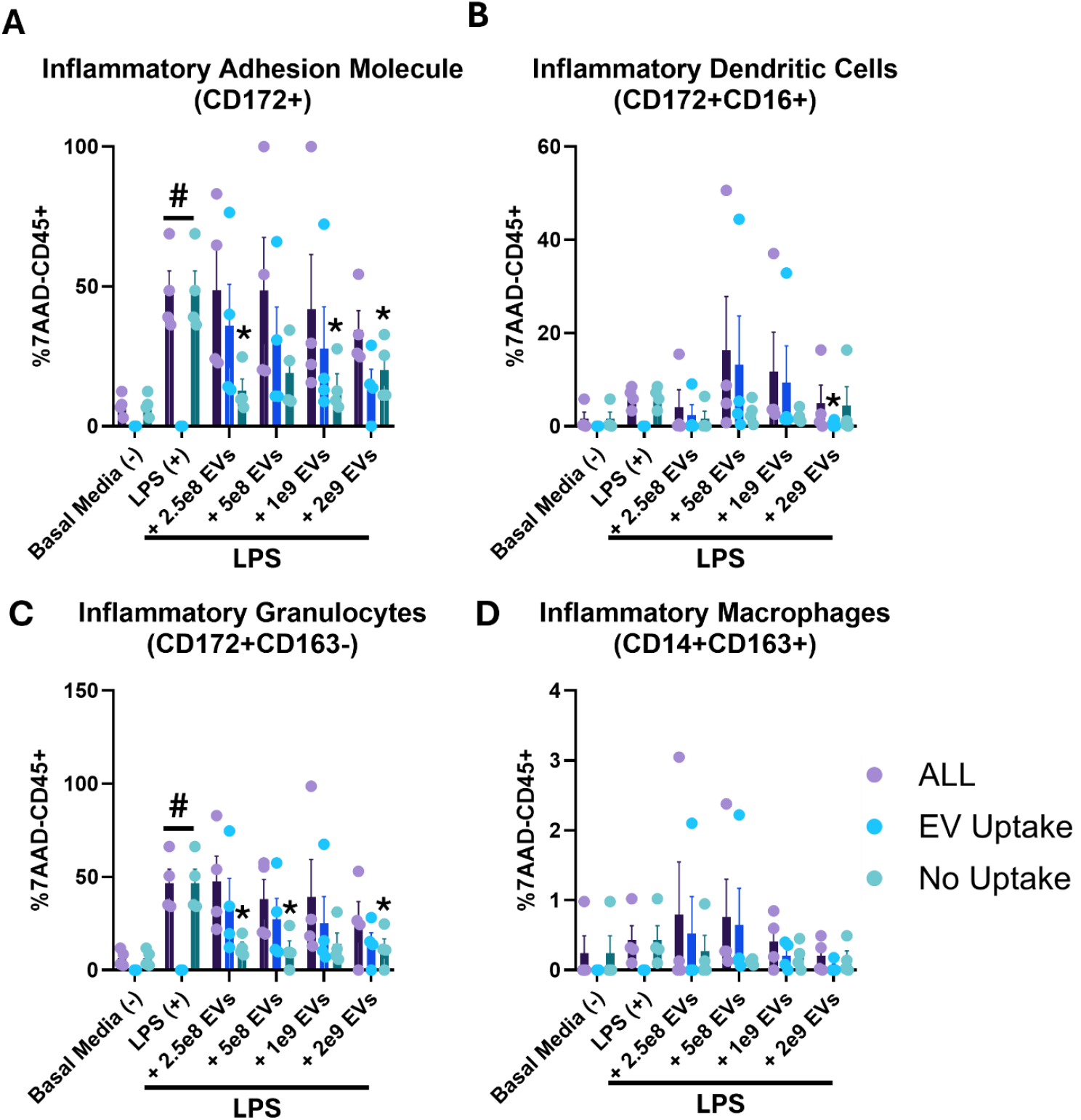
MSC-EVs Attenuate LPS-Induced Inflammatory Shifts in CD172+ Cells, Dendritic Cells, Granulocytes, and Macrophages. Relative porcine PBMC (%7AAD-/CD45+) surface marker expression of A) inflammatory adhesion molecule (CD172+) and inflammatory B) dendritic cells (CD172+CD16+), C) granulocytes (CD172+CD163-) and D) macrophages (CD14+CD163+) following co-culture with LPS and treated with either +2.5e8 MSC-EVs, +5e8 MSC-EVs, +1e9 MSC-EVs, or +2e9 MSC-EVs against basal media (-) control. Data presented as Mean ± SEM. n=biological triplicates, technical duplicates. *p<0.05 vs. LPS. #p<0.05 vs. Basal Media (-).

### Inhibition of LPS-Induced Inflammatory Transcript Levels by MSC-EVs

To assess the transcriptional effects of MSC-EV treatment on LPS-stimulated PBMCs, we conducted targeted qPCR to quantify key inflammatory transcripts. LPS induced a significant increase in IFNα expression that was completely attenuated by MSC-EVs, regardless of concentration (**Figure 3A**). Likewise, IL-1α levels were significantly reduced by treatment with each concentration of MSC-EVs (**Figure 3B**). LPS-induced IL-1β expression decreased with MSC-EV concentrations of 2.5e8, 5e8, and 1e9, but showed a notable increase when 2e9 MSC-EVs were added (**Figure 3C**). IL-6 transcripts followed a similar MSC-EV concentration-independent pattern as IFNα (**Figure 3D**), while LPS-induced expression of IL-8 was attenuated by each concentration of MSC-EVS (**Figure 3E**). Finally, LPS-induced elevations in TNFα expression were also attenuated by MSC-EVs, with the most prominent effects observed in PBMCs treated with intermediate concentrations (5e8 and 1e9; **Figure 3F**). Overall, these observations indicate that MSC-derived EVs can differentially modulate inflammatory gene expression in LPS-activated PBMCs, with distinct concentration-specific effects across multiple cytokines (**Figure S6A and Table S5**). Such findings underscore the potential of EVs as key immunomodulatory regulators of LPS-induced immune responses, commonly seen as “sterile sepsis” in PCAS.

**Figure 3:**
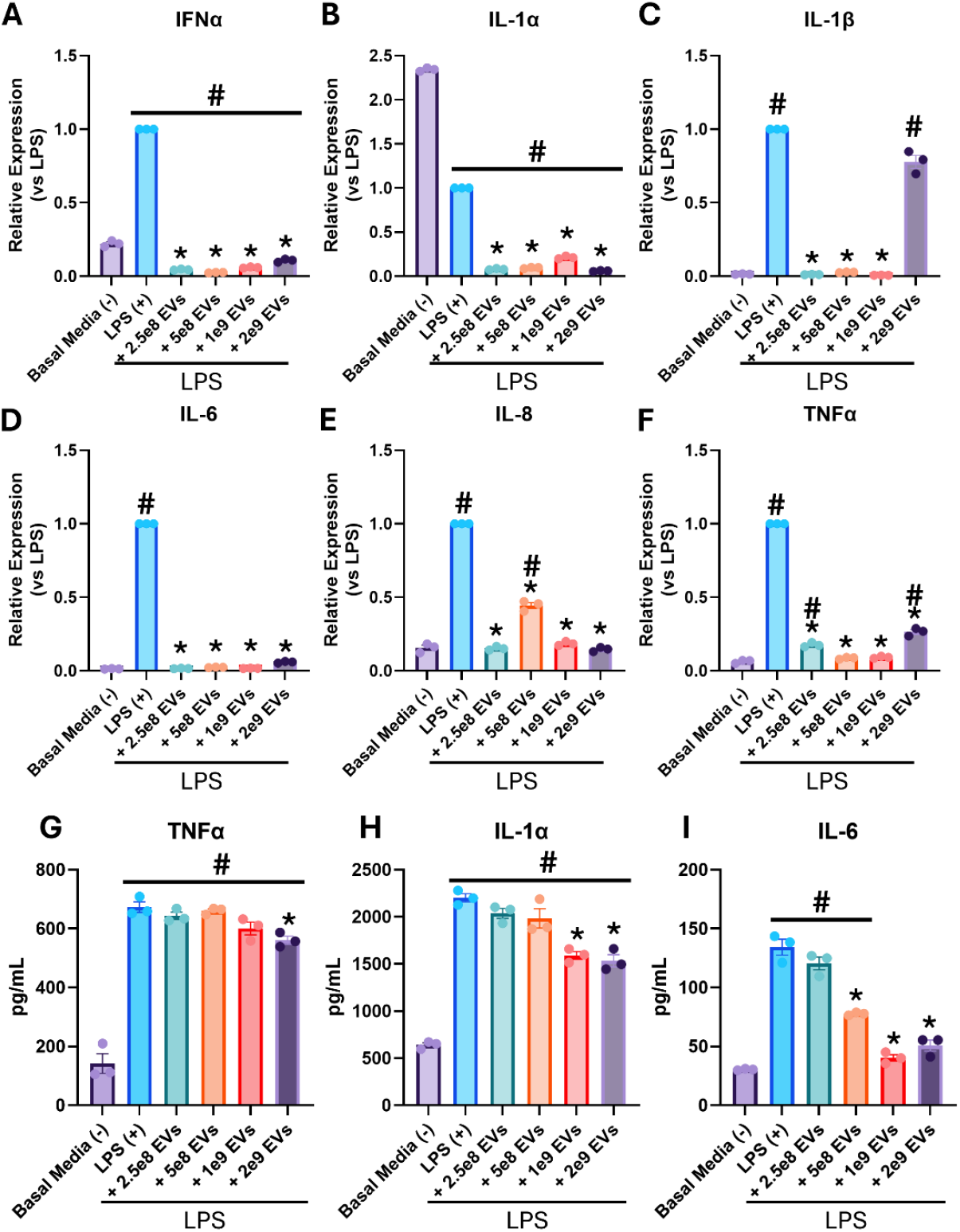
MSC-EVs Reduce Inflammatory Gene Expression and Cytokine Secretion in Porcine PBMCs Activated by LPS. Relative PBMC inflammatory transcript expression of A) IFNα, B) IL-1α, C) IL-1β, D) IL-6, E) IL-8, and F) TNFα at 24-hr post-activation assay with LPS and treated with either +2.5e8 MSC-EVs, +5e8 MSC-EVs, +1e9 MSC-EVs, or +2e9 MSC-EVs vs. basal media (-) control. Housekeeping gene = β2M. Conditioned media levels of PBMC-secreted inflammatory cytokines, G) TNFα, H) IL-1β, and I) IL-6 following a 24hr activation assay with LPS and treated with +2.5e8 MSC-EVs, +5e8 MSC-EVs, +1e9 MSC-EVs, or +2e9 MSC-EVs against basal media (-) control. Data presented as Mean ± SEM. n=biological triplicates, technical duplicates. *p<0.05 vs. LPS. #p<0.05 vs. Basal Media (-). The horizontal line indicates that all bars below are significant.

### Suppression of LPS-Stimulated Inflammatory Cytokine Release by MSC-EVs

To further examine the immunomodulatory capacity of MSC-EVs on LPS-stimulated PBMCs, we quantified TNFα, IL-1α, IL-6, and IL-10 levels by ELISA to assess potential concentration-dependent effects (**Figure S7A and Table S6**). TNFα concentrations remained largely unchanged relative to the LPS-only control, except for a significant reduction at the highest concentration of MSC-EVs (2e9; **Figure 3G**).

In contrast, IL-1α exhibited a concentration-dependent decline following MSC-EV treatment, with significant decreases observed at 1e9 and 2e9 MSC-EVs (**Figure 3H**). The most pronounced response to MSC-EV treatment was observed with respect to IL-6 levels, where MSC-EV concentrations of 5e8, 1e9, and 2e9 produced substantial reductions compared to LPS-stimulation alone (**Figure 3I**). These results collectively indicate that MSC-EVs exert robust immunomodulatory effects on pro-inflammatory cytokine release from PBMCs, with higher MSC-EV concentrations attenuating LPS-induced production of TNFα, IL-1α, and IL-6.

### Modulation of mtDNA-Triggered Inflammatory Cell Phenotype by MSC-EVs

While LPS is thought to provoke delayed immune cell activation in PCAS (∼24-48 hours post-ROSC), our laboratory has recently shown that circulating levels of cell-free mtDNA increase within the first few hours after resuscitation and can trigger early activation of PBMCs following cytoplasmic mtDNA internalization (7, 8, 29, 30). Thus, we were interested in evaluating the effects of MSC-EVs on naïve PBMCs activated by internalized mtDNA (via transfection with Lipofectamine 2000; TR) to mimic early immune cell activation in PCAS. First, flow cytometric analyses were conducted across a range of immune cell subsets and, in contrast to the selective effects observed in LPS-activated PBMCs (**Figure 2**), DiD-labeled MSC-EVs robustly inhibited mtDNA-induced phenotypic shifts in all inflammatory subpopulations examined (**Figure 4**). Specifically, the highest MSC-EV concentration (2e9) resulted in a statistically significant reduction in CD172+ cells (**Figure 4A and S5A**). Similarly, the number of inflammatory dendritic cells was significantly reduced at all MSC-EV concentrations, with the largest effect observed with 2e9 MSC-EVs (**Figure 4B and S5B**). Inflammatory granulocytes mirrored this trend, as did cells expressing inflammatory adhesion molecule CD172, as each subpopulation was significantly reduced by treatment with 2e9 MSC-EVs (**Figure 4C and S5C**). In contrast, inflammatory macrophages were significantly reduced by all MSC-EV concentrations except the lowest (2.5e8) (**Figure 4D and S5D**). Notably, these immunosuppressive effects were observed in both MSC-EV-uptake and no-EV-uptake cell populations and occurred in the absence of cell death (**Figure S4B**), underscoring a broad immunomodulatory capacity of MSC-EVs that was not dependent on cell uptake (**Table S4)**. Collectively, these findings highlight concentration-dependent anti-inflammatory properties of MSC-EVs by demonstrating their ability to attenuate mtDNA-induced shifts of naïve PBMCs towards pro-inflammatory phenotypes.

**Figure 4:**
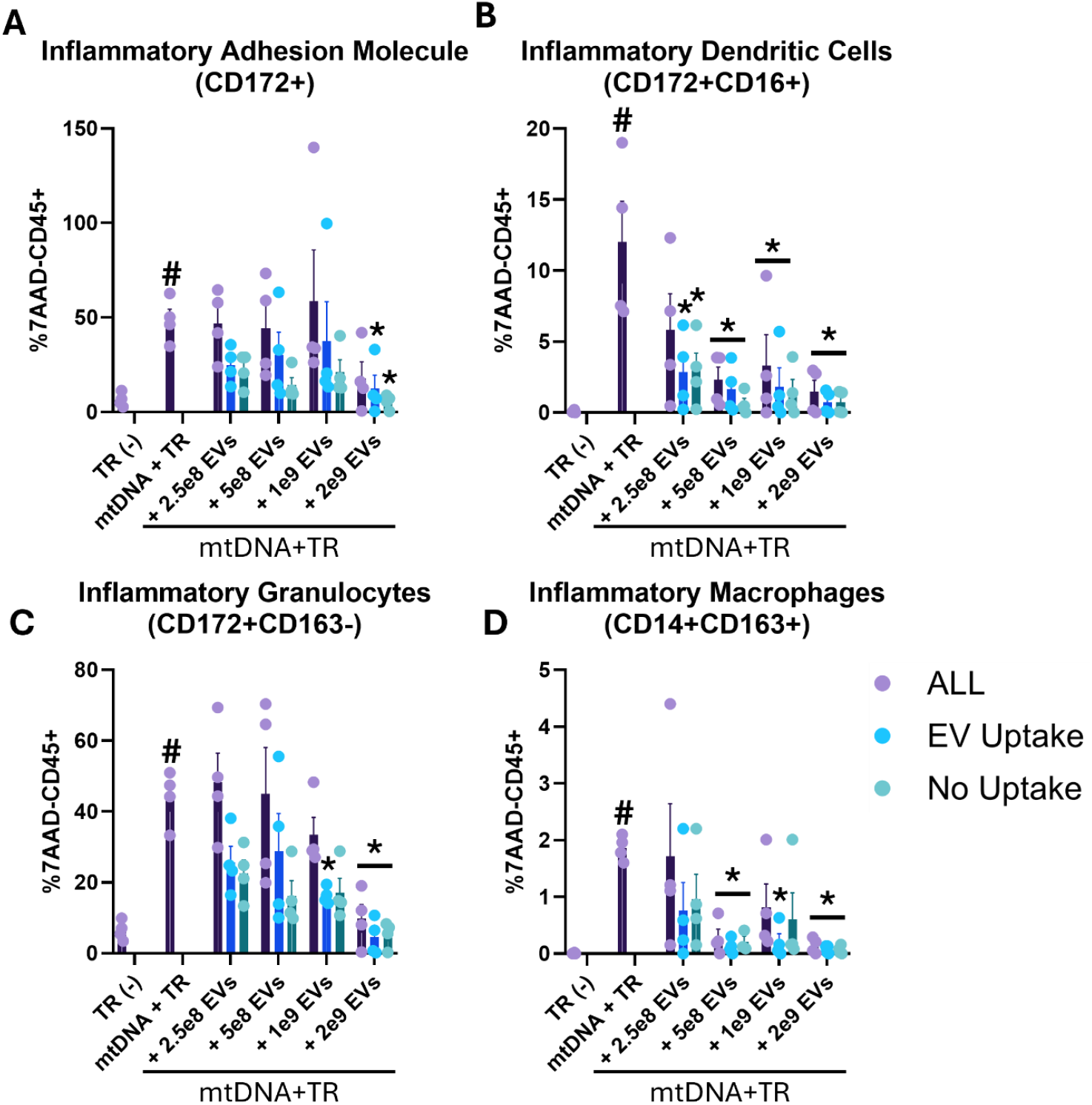
MSC-EVs Attenuate mtDNA-Induced Inflammatory Shifts in CD172+ Cells, Dendritic Cells, Granulocytes, and Macrophages. Relative porcine PBMC (%7AAD-/CD45+) surface marker expression of A) inflammatory adhesion molecule (CD172+) and inflammatory B) dendritic cells (CD172+CD16+), C) granulocytes (CD172+CD163-) and D) macrophages (CD14+CD163+) following co-culture with mtDNA+TR and treated with either +2.5e8 MSC-EVs, +5e8 MSC-EVs, +1e9 MSC-EVs, or +2e9 MSC-EVs against transfection reagent (TR; negative control). Data presented as Mean ± SEM. n=biological triplicates, technical duplicates. *p<0.05 vs. mtDNA+TR. #p<0.05 vs. TR (-).

### Inhibition of mtDNA-Driven Inflammatory Gene Expression by MSC-EVs

To evaluate the transcriptional impact of MSC-EV treatment in mtDNA-stimulated PBMCs, we performed qPCR to evaluate expression of key inflammatory transcripts. IFNα expression increased in cells transfected with mtDNA but was significantly decreased by all MSC-EV concentrations (**Figure 5A**). IL-1α followed a similar trend, with the highest expression in mtDNA-only cultures and a concentration-independent reduction in response to MSC-EVs (**Figure 5B**). mtDNA-induced IL-1β expression also displayed a pronounced decrease in response to MSC-EVs, as all MSC-EV concentrations led to a marked decrease in transcripts (**Figure 5C**). In contrast, IL-6 expression did not uniformly diminish across the treated samples; however, 1e9 and 2e9 MSC-EVs significantly decreased IL-6 expression compared with the mtDNA + TR group (**Figure 5D**). IL-8 was notably reduced in all MSC-EV conditions, with the lowest levels observed at the 2.5e8 and 1e9 concentrations (**Figure 5E**). Lastly, TNFα expression displayed a concentration-dependent decrease following MSC-EV treatment, though the 2.5e8 dose interestingly yielded higher expression than the mtDNA-positive control (**Figure 5F**). Overall, these results reveal a concentration-dependent immunomodulatory influence of MSC-EVs in mtDNA-activated PBMCs (**Figure S6B and Table S5**), albeit with variable potency across distinct cytokine targets.

**Figure 5:**
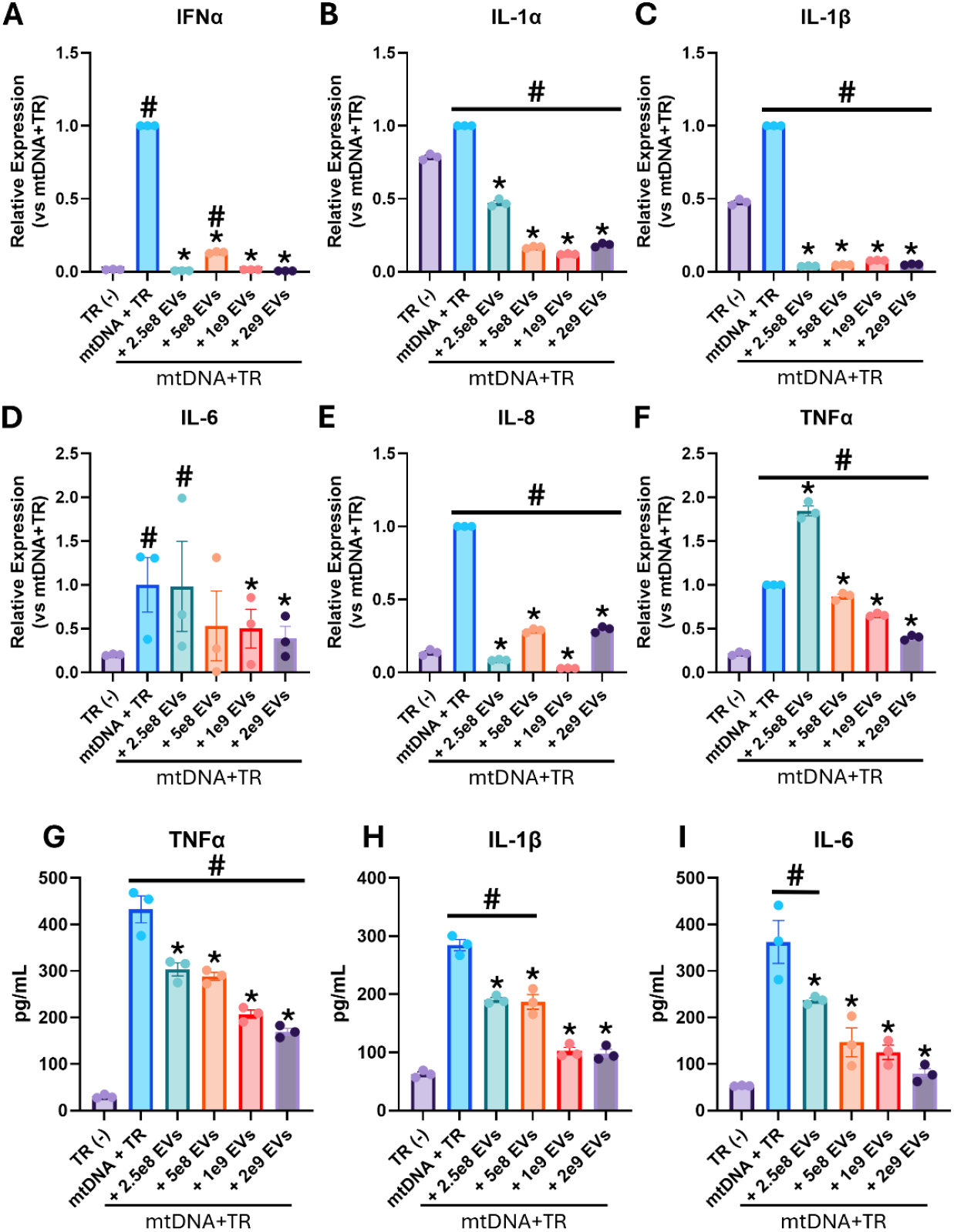
MSC-EVs Reduce Inflammatory Gene Expression and Cytokine Secretion in Porcine PBMCs Activated by mtDNA. Relative PBMC inflammatory transcript expression of A) IFNα, B) IL-1α, C) IL-1β, D) IL-6, E) IL-8, and F) TNF-α at 24-hr post-activation assay with mtDNA+TR and treated with either +2.5e8 MSC-EVs, +5e8 MSC-EVs, +1e9 MSC-EVs, or +2e9 MSC-EVs vs. transfection reagent (TR; negative control). Housekeeping gene = B2M. Conditioned media levels of PBMC-secreted inflammatory cytokines, G) TNFα, H) IL-1β, and I) IL-6 following a 24hr activation assay with mtDNA+TR and treated with +2.5e8 MSC-EVs, +5e8 MSC-EVs, +1e9 MSC-EVs, or +2e9 MSC-EVs against transfection reagent (TR; negative control). Data presented as Mean ± SEM. n=biological triplicates, technical duplicates. *p<0.05 vs. mtDNA+TR. #p<0.05 vs. TR (-).

### Attenuated mtDNA-Induced Inflammatory Cytokine Production by MSC-EVs

To investigate the concentration-dependent immunomodulatory effects of MSC-EVs in mtDNA-stimulated PBMCs, we performed ELISA to quantify TNFα, IL-1β, IL-6, and IL-10 (**Figure S7B and Table S6**). TNFα levels increased following mtDNA-stimulation but were significantly reduced by treatment with increasing concentrations of MSC-EVs (**Figure 5G**). mtDNA-induced IL-1β secretion was also attenuated across all MSC-EV concentrations, with the two highest concentrations (1e9 and 2e9 MSC-EVs) reducing IL-1β levels to values comparable to the negative control (**Figure 5H**). Similarly, IL-6 displayed a clear concentration-dependent decline following MSC-EV treatment, achieving statistical significance at all MSC-EV concentrations (**Figure 5I**). Collectively, these findings highlight a nuanced, concentration-dependent modulatory role of MSC-EVs in curbing mtDNA-induced inflammatory responses, underscoring their potential ability to ameliorate mtDNA-driven cytokine production in PCAS.

### Reduction of ROS/RNS Generation in LPS- and mtDNA-Activated Cells via MSC-EVs

Monocytes and neutrophils generate reactive oxygen species (ROS) via myeloperoxidase, which can modulate neighboring cells and drive inflammatory processes associated with PCAS and IRI (31). To explore whether MSC-EV treatment attenuates ROS and reactive nitrogen species (RNS) production, we quantified ROS+RNS levels in the conditioned media of cells stimulated by LPS or mtDNA. In LPS-stimulated samples, MSC-EV administration produced a concentration-dependent reduction in ROS/RNS, with MSC-EV concentrations above 5e8 significantly lowering levels below those of the negative control (**Figure 6A and S8; Table S6**). Similarly, in mtDNA-stimulated PBMCs, 5e8 and 2e9 MSC-EV concentrations significantly reduced ROS/RNS (**Figure 6B and S8; Table S6**). These findings support a protective role for MSC-EVs in mitigating oxidative stress under inflammatory conditions relevant to PCAS.

**Figure 6:**
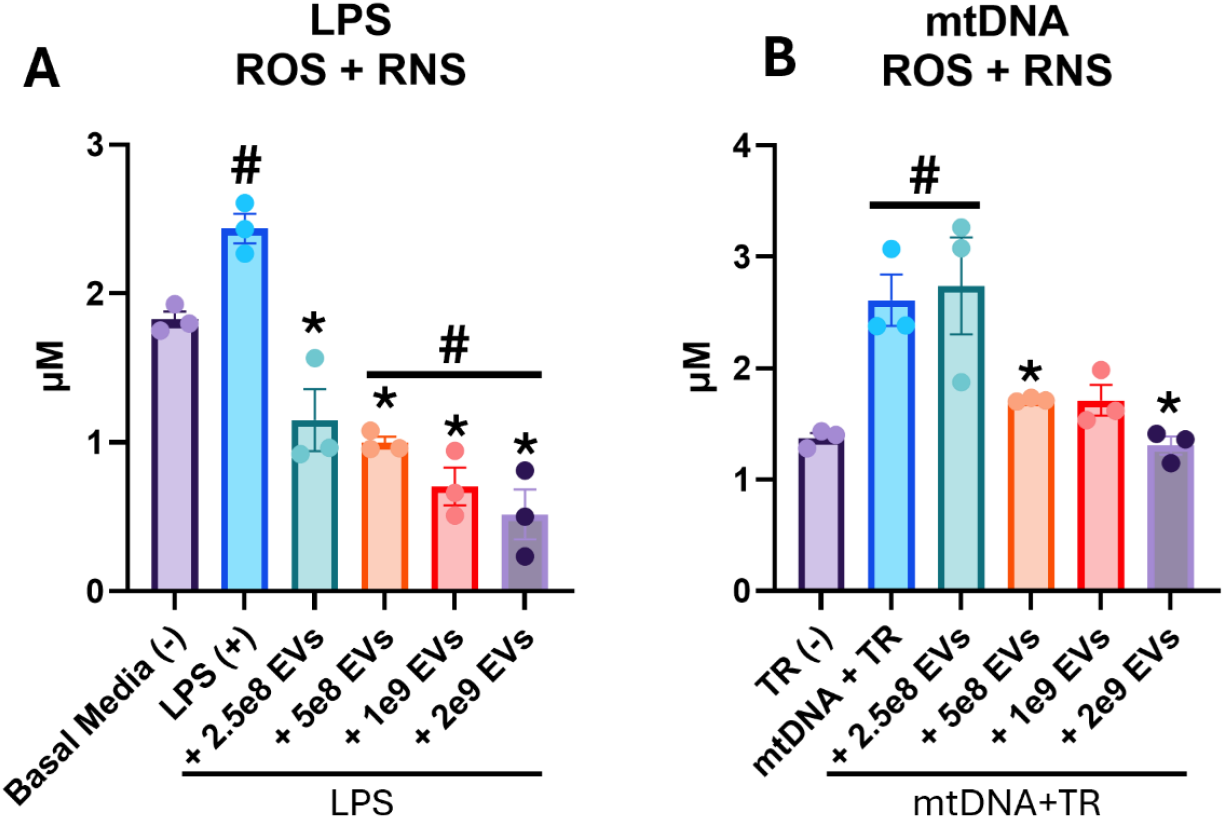
MSC-EVs Attenuate LPS- and mtDNA-Induced ROS and RNS Production. Conditioned media levels of reactive oxygen and nitrogen species (ROS/RNS) following a 24hr post-activation assay with A) LPS or B) mtDNA+TR and treated with either +2.5e8 MSC-EVs, +5e8 MSC-EVs, +1e9 MSC-EVs, or +2e9 MSC-EVs against transfection reagent (TR; negative control). Data presented as Mean ± SEM. n=biological triplicates, technical duplicates. *p<0.05 vs. mtDNA+TR. #p<0.05 vs. Basal Media (-) or TR (-).

## DISCUSSION

In this study, we explored how MSC-EVs modulate inflammatory responses in naïve porcine PBMCs activated with either LPS or mtDNA to mimic immune cell activation in PCAS. Notably, MSC-EV treatment attenuated LPS-induced inflammatory granulocyte activation, downregulated multiple inflammatory cytokine transcripts, suppressed pro-inflammatory cytokine secretion, and mitigated oxidative stress. Following mtDNA stimulation, MSC-EVs inhibited immune activation in dendritic cells, granulocytes, and macrophages, while also reducing expression and release of key inflammatory cytokines as well as ROS/RNS levels. These observations support the overarching conclusion that MSC-EVs exhibit potent immunomodulatory properties and are capable of modulating multiple pro-inflammatory parameters in immune cells activated by PCAS-relevant stimuli.

The present findings add to an accumulating body of evidence that MSCs may exert much of their therapeutic benefit via paracrine mechanisms, chiefly mediated by EVs (11, 32). EVs, which include exosomes and microvesicles, transport bioactive cargo such as proteins, lipids, and regulatory RNAs (e.g., miRNAs) that can profoundly influence recipient cell function (33-35). In the context of IRI and other pro-inflammatory conditions, MSC-EVs have shown promise in diminishing infarct size, enhancing angiogenesis, and limiting fibrotic remodeling (19). The current work complements these findings by demonstrating MSC-EVs’ capacity to modulate immune cell phenotypes and inflammatory signaling cascades following stimulation with LPS or mtDNA. This immunomodulatory effect is particularly relevant to PCAS, where systemic inflammation triggered, at least in part, by LPS- and mtDNA-induced immune cell activation exacerbates end-organ damage and worsens patient outcomes (1).

Our findings build on previous reports showing that MSC-EVs can attenuate sterile inflammatory responses triggered by high-mobility group box 1 (HMGB1) and other damage-associated molecular patterns (DAMPs) (36). Here, we present new evidence that MSC-EVs significantly inhibit LPS-driven immune activation at both the cellular and transcript levels. Critically, MSC-EV concentrations above 5e8 dramatically reduced ROS/RNS generation in LPS-stimulated PBMCs, suggesting that oxidative stress— known to contribute to PCAS pathophysiology—may be tempered by MSC-EV treatment. Perhaps even more striking is the suppression of mtDNA-induced inflammatory phenotypes across multiple immune cell populations, including inflammatory dendritic cells, granulocytes, and macrophages, supporting prior observations that MSC-EVs can mitigate TLR9-mediated responses (via which mtDNA may exert its inflammatory activation (37)). This comprehensive inhibition of mtDNA-induced cytokine production positions MSC-EVs as a promising intervention against sterile inflammatory triggers relevant to PCAS and other conditions characterized by IRI.

## Limitations

Despite these encouraging results, several experimental limitations warrant discussion. First, our *in vitro* study employs a porcine PBMC model, which, while valuable for translational work, may differ from human immune cell responses. Second, the mechanistic details of how specific MSC-EV cargo (e.g., miRNAs, proteins) modulate cytokine transcripts and oxidative stress remain incompletely characterized. Elucidating these mechanisms would aid in refining EV-based therapies for targeted immunomodulation. Third, MSC-EV concentration-dependent effects observed in the present study suggest a therapeutic window, but large-scale dosing studies and *in vivo* validation in clinically relevant models (e.g., porcine PCAS models) are necessary to fully characterize the safety and efficacy of MSC-EVs (38). Finally, we have not examined the long-term stability or biodistribution of MSC-EVs *in vivo*, factors that are crucial for eventual therapeutic application (39).

## Conclusions

The results of the present study demonstrate the potent immunomodulatory properties of MSC-EVs against LPS- and mtDNA-induced inflammation in porcine PBMCs. By attenuating pro-inflammatory cytokine expression, reducing immune cell activation, and mitigating ROS/RNS production, MSC-EVs show promise as a therapeutic strategy for PCAS and other acute inflammatory conditions. These results strengthen the growing consensus that MSC-EVs act as key paracrine effectors in MSC-based therapies (11). Future investigation focusing on cargo-specific mechanisms, optimal dosing regimens, and translational *in vivo* studies will be critical in moving toward effective EV-based therapeutics for PCAS as well as other pro-inflammatory disease states involving IRI.

## GLOSSARY

PCAS: Post-Cardiac Arrest Syndrome
SCA: Sudden Cardiac Arrest
ROSC: Return of Spontaneous Circulation
IRI: Ischemia Reperfusion Injury
LPS: Lipopolysaccharide
mtDNA: Mitochondrial DNA
PBMC: Peripheral Blood Mononuclear Cell
MSC: Mesenchymal Stem Cell
MSC-EV: Mesenchymal Stem Cell - Extracellular Vesicle
TR: Transfection Reagent
FBS: Fetal Bovine Serum
CM: Conditioned Media
ROS: Reactive Oxygen Species
RNS: Reactive Nitrogen Species

## DATA AVAILABILITY

All summarized data and supporting materials have been provided with the published article. Raw data is available from the corresponding author upon reasonable request.

## SUPPLEMENTAL MATERIAL

See Supplemental Materials for expanded methods, figures, and tables.

## ACKNOWLEDGMENTS

This project was made possible by support from the University at Buffalo, NY core facilities and the Roswell Park Comprehensive Cancer Center Flow and Image Cytometry Shared Resource. These studies could not have been completed without the excellent technical support of Maggie Vogel-Cryan, LVT, and Beth Palka. We acknowledge the use of BioRender for figure design.

## REFERENCES

1. Neumar RW, Nolan JP, Adrie C, Aibiki M, Berg RA, Böttiger BW, et al. Post-cardiac arrest syndrome: epidemiology, pathophysiology, treatment, and prognostication. A consensus statement from the International Liaison Committee on Resuscitation (American Heart Association, Australian and New Zealand Council on Resuscitation, European Resuscitation Council, Heart and Stroke Foundation of Canada, InterAmerican Heart Foundation, Resuscitation Council of Asia, and the Resuscitation Council of Southern Africa); the American Heart Association Emergency Cardiovascular Care Committee; the Council on Cardiovascular Surgery and Anesthesia; the Council on Cardiopulmonary, Perioperative, and Critical Care; the Council on Clinical Cardiology; and the Stroke Council. Circulation. 2008;118(23):2452–83.

2. Weil BR, Allen SE, Barbaccia T, Wong K, Beaver AM, Slabinski EA, et al. Preclinical evaluation of triiodothyronine nanoparticles as a novel therapeutic intervention for resuscitation from cardiac arrest. Resuscitation. 2023;186:109735.

3. Adrie C, Adib-Conquy M, Laurent I, Monchi M, Vinsonneau C, Fitting C, et al. Successful cardiopulmonary resuscitation after cardiac arrest as a “sepsis-like” syndrome. Circulation. 2002;106(5):562–8.

4. Zhang Q, Raoof M, Chen Y, Sumi Y, Sursal T, Junger W, et al. Circulating mitochondrial DAMPs cause inflammatory responses to injury. Nature. 2010;464(7285):104–7.

5. Krysko DV, Agostinis P, Krysko O, Garg AD, Bachert C, Lambrecht BN, et al. Emerging role of damage-associated molecular patterns derived from mitochondria in inflammation. Trends Immunol. 2011;32(4):157–64.

6. Cucinotta DM, Rolland T, Zahra S, Weil BR. Abstract P3063: Selective Release Of Mitochondrial Dna After Brief Myocardial Ischemia. Circulation Research. 2023;133(Suppl_1):AP3063–AP.

7. Rolland T, Cucinotta D, Zahra S, Weil BR. Abstract 171: Hyperoxia Potentiates Pro-Inflammatory Immune Cell Activation in vitro. Circulation. 2023;148(Suppl_1):A171–A.

8. Zahra S, Rolland TJ, Cucinotta D, Hudson ER, Young R, Weil B. Circulating Extracellular Vesicle-Mediated Immune Activation After Resuscitation From Cardiac Arrest in Swine. Physiology. 2024;39(S1):1310.

9. Nolan JP, Soar J, Cariou A, Cronberg T, Moulaert VR, Deakin CD, et al. European Resuscitation Council and European Society of Intensive Care Medicine Guidelines for Post-resuscitation Care 2015: Section 5 of the European Resuscitation Council Guidelines for Resuscitation 2015. Resuscitation. 2015;95:202–22.

10. Chen J, Li Y, Wang L, Zhang Z, Lu D, Lu M, et al. Therapeutic benefit of intravenous administration of bone marrow stromal cells after cerebral ischemia in rats. Stroke. 2001;32(4):1005–11.

11. Gnecchi M, Zhang Z, Ni A, Dzau VJ. Paracrine mechanisms in adult stem cell signaling and therapy. Circ Res. 2008;103(11):1204–19.

12. Imberti B, Morigi M, Tomasoni S, Rota C, Corna D, Longaretti L, et al. Insulin-like growth factor-1 sustains stem cell mediated renal repair. J Am Soc Nephrol. 2007;18(11):2921–8.

13. Tajlil A, Zel R, Wadhwani A, Nsumbu D, Young R, Weil BR. Abstract 107: Systemic Allogeneic Mesenchymal Stem Cell Administration Does Not Attenuate Early Post-resuscitation Brain Injury Or Inflammation In A Porcine Model Of Cardiac Arrest. Circulation. 2022;146(Suppl_1):A107–A.

14. Chong SY, Lee CK, Huang C, Ou YH, Charles CJ, Richards AM, et al. Extracellular Vesicles in Cardiovascular Diseases: Alternative Biomarker Sources, Therapeutic Agents, and Drug Delivery Carriers. Int J Mol Sci. 2019;20(13).

15. Saludas L, Oliveira CC, Roncal C, Ruiz-Villalba A, Prósper F, Garbayo E, et al. Extracellular Vesicle-Based Therapeutics for Heart Repair. Nanomaterials (Basel). 2021;11(3).

16. Mackie AR, Klyachko E, Thorne T, Schultz KM, Millay M, Ito A, et al. Sonic hedgehog-modified human CD34+ cells preserve cardiac function after acute myocardial infarction. Circ Res. 2012;111(3):312–21.

17. Wang Y, Zhang L, Li Y, Chen L, Wang X, Guo W, et al. Exosomes/microvesicles from induced pluripotent stem cells deliver cardioprotective miRNAs and prevent cardiomyocyte apoptosis in the ischemic myocardium. Int J Cardiol. 2015;192:61–9.

18. Liu L, Jin X, Hu CF, Li R, Zhou Z, Shen CX. Exosomes Derived from Mesenchymal Stem Cells Rescue Myocardial Ischaemia/Reperfusion Injury by Inducing Cardiomyocyte Autophagy Via AMPK and Akt Pathways. Cell Physiol Biochem. 2017;43(1):52–68.

19. Arslan F, Lai RC, Smeets MB, Akeroyd L, Choo A, Aguor EN, et al. Mesenchymal stem cell-derived exosomes increase ATP levels, decrease oxidative stress and activate PI3K/Akt pathway to enhance myocardial viability and prevent adverse remodeling after myocardial ischemia/reperfusion injury. Stem Cell Res. 2013;10(3):301–12.

20. Aguiar Koga BA, Fernandes LA, Fratini P, Sogayar MC, Carreira ACO. Role of MSC-derived small extracellular vesicles in tissue repair and regeneration. Front Cell Dev Biol. 2022;10:1047094.

21. Lai RC, Arslan F, Lee MM, Sze NS, Choo A, Chen TS, et al. Exosome secreted by MSC reduces myocardial ischemia/reperfusion injury. Stem Cell Res. 2010;4(3):214–22.

22. Lai RC, Chen TS, Lim SK. Mesenchymal stem cell exosome: a novel stem cell-based therapy for cardiovascular disease. Regen Med. 2011;6(4):481–92.

23. Yang S, Li J, Tang M, Gao X, Liu W, Wei S. Mesenchymal Stem Cell-Derived Exosomes in Cardioprotection: A Novel Application to Prevent Myocardial Injury. Rev Cardiovasc Med. 2022;23(9):310.

24. Wang X, Huang W, Liu G, Cai W, Millard RW, Wang Y, et al. Cardiomyocytes mediate anti-angiogenesis in type 2 diabetic rats through the exosomal transfer of miR-320 into endothelial cells. J Mol Cell Cardiol. 2014;74:139–50.

25. Racchetti G, Meldolesi J. Extracellular Vesicles of Mesenchymal Stem Cells: Therapeutic Properties Discovered with Extraordinary Success. Biomedicines. 2021;9(6).

26. Milano G, Biemmi V, Lazzarini E, Balbi C, Ciullo A, Bolis S, et al. Intravenous administration of cardiac progenitor cell-derived exosomes protects against doxorubicin/trastuzumab-induced cardiac toxicity. Cardiovasc Res. 2020;116(2):383–92.

27. Zhang Y, Mei H, Chang X, Chen F, Zhu Y, Han X. Adipocyte-derived microvesicles from obese mice induce M1 macrophage phenotype through secreted miR-155. J Mol Cell Biol. 2016;8(6):505–17.

28. Hartjes TA, Mytnyk S, Jenster GW, van Steijn V, van Royen ME. Extracellular Vesicle Quantification and Characterization: Common Methods and Emerging Approaches. Bioengineering (Basel). 2019;6(1).

29. West AP, Khoury-Hanold W, Staron M, Tal MC, Pineda CM, Lang SM, et al. Mitochondrial DNA stress primes the antiviral innate immune response. Nature. 2015;520(7548):553–7.

30. Paludan SR. Activation and regulation of DNA-driven immune responses. Microbiol Mol Biol Rev. 2015;79(2):225–41.

31. Kornfeld OS, Hwang S, Disatnik MH, Chen CH, Qvit N, Mochly-Rosen D. Mitochondrial reactive oxygen species at the heart of the matter: new therapeutic approaches for cardiovascular diseases. Circ Res. 2015;116(11):1783–99.

32. Rani S, Ryan AE, Griffin MD, Ritter T. Mesenchymal Stem Cell-derived Extracellular Vesicles: Toward Cell-free Therapeutic Applications. Mol Ther. 2015;23(5):812–23.

33. Chen TS, Lai RC, Lee MM, Choo AB, Lee CN, Lim SK. Mesenchymal stem cell secretes microparticles enriched in pre-microRNAs. Nucleic Acids Res. 2010;38(1):215–24.

34. Rautou PE, Vion AC, Amabile N, Chironi G, Simon A, Tedgui A, et al. Microparticles, vascular function, and atherothrombosis. Circ Res. 2011;109(5):593–606.

35. Hergenreider E, Heydt S, Tréguer K, Boettger T, Horrevoets AJ, Zeiher AM, et al. Atheroprotective communication between endothelial cells and smooth muscle cells through miRNAs. Nat Cell Biol. 2012;14(3):249–56.

36. Homma K, Bazhanov N, Hashimoto K, Shimizu M, Heathman T, Hao Q, et al. Mesenchymal stem cell-derived exosomes for treatment of sepsis. Front Immunol. 2023;14:1136964.

37. West AP, Shadel GS, Ghosh S. Mitochondria in innate immune responses. Nat Rev Immunol. 2011;11(6):389–402.

38. Chen L, Wang Y, Pan Y, Zhang L, Shen C, Qin G, et al. Cardiac progenitor-derived exosomes protect ischemic myocardium from acute ischemia/reperfusion injury. Biochem Biophys Res Commun. 2013;431(3):566–71.

39. Lai RC, Yeo RWY, Tan SS, Zhang B, Yin Y, Sze NSK, et al., editors. Mesenchymal Stem Cell Exosomes: The Future MSC-Based Therapy? 2013.

40. Roth S, Rottach A, Lotz-Havla AS, Laux V, Muschaweckh A, Gersting SW, et al. Rad50-CARD9 interactions link cytosolic DNA sensing to IL-1β production. Nat Immunol. 2014;15(6):538–45.

41. Ezquerra A, Revilla C, Alvarez B, Pérez C, Alonso F, Domínguez J. Porcine myelomonocytic markers and cell populations. Dev Comp Immunol. 2009;33(3):284–98.

42. Grandoni F, Scatà MC, Martucciello A, De Carlo E, De Matteis G, Hussen J. Comprehensive phenotyping of peripheral blood monocytes in healthy bovine. Cytometry A. 2022;101(2):122–30.

43. Piriou-Guzylack L, Salmon H. Membrane markers of the immune cells in swine: an update. Vet Res. 2008;39(6):54.

44. Álvarez B, Revilla C, Poderoso T, Ezquerra A, Domínguez J. Porcine Macrophage Markers and Populations: An Update. Cells. 2023;12(16).

45. Krajnik A, Nimmer E, Brazzo JA, 3rd, Biber JC, Drewes R, Tumenbayar BI, et al. Survivin regulates intracellular stiffness and extracellular matrix production in vascular smooth muscle cells. APL Bioeng. 2023;7(4):046104.

46. Heidari R, Taheri V, Rahimi HR, Shirazi Yeganeh B, Niknahad H, Najibi A. Sulfasalazine-induced renal injury in rats and the protective role of thiol-reductants. Ren Fail. 2016;38(1):137–41.

47. Zhang F. WK, Wu X., Xin C., Zhou M., Lei J., Chen J. Punicalagin Alleviates Brain Injury and Inflammatory Responses, and Regulates HO-1/Nrf-2/ARE Signaling in Rats after Experimental Intracerebral Haemorrhage. Trop J Pharm Res. 2020(19):727–37.

