## Supplemental Materials + Figures + Tables for "Mesenchymal Stem Cell-Derived Extracellular Vesicles Mitigate Immune Cell Activation in an In Vitro Model of Post-Resuscitation Inflammation"

#### Expanded Methods

Ethics Statement

Investigators were blinded to conditions (e.g., mtDNA vs. mtDNA + MSC-EVs) while conducting analytical protocols and analyzing data. All experimental procedures and protocols conformed to institutional guidelines for the care and use of animals in research and were approved by the State University of New York at Buffalo Institutional Animal Care and Use Committee.

Bone Marrow - MSC Isolation and Culture

Bone marrow derived mesenchymal stem cells (MSCs) were isolated from the bone marrow of healthy White Yorkshire x Landrace swine (Oak Hill Genetics, Ewing, IL) and cultivated in Advanced DMEM (Gibco #12491) supplemented with Glutamax (Invitrogen #35050-061), 10% Fetal Bovine Serum (FBS) and penicillin/streptomycin. Approximately 16 mL of whole bone marrow was collected and added to Ficoll CPT Vacutainers (BD, Ref 362761), followed by centrifugation at 2,500rpm for 25 minutes at 25°C. The upper plasma layer was discarded and the buffy layer, containing MSCs, added to 40ml PBS/1% BSA. Following a 15-minute centrifugation at 300xg, the pellet was resuspended in media and plated in a 100mm dish (P0). The next day, when the cells were attached, they were washed 3X with PBS, to minimize fungal growth, and new media was added. Wash/feed was continued every other day for 7-10 days or until cells were dense enough for passage with trypsin. PBS washes were not included with feeding beyond passage 0. MSCs were cryopreserved in ADMEM culture media (specifications above), 1x B-27 Supplement (Gibco, Ref 17504044), and 10% DMSO slowly to -80°C before being transferred to liquid nitrogen for long term storage.

MSC-EV Isolation and Characterization

MSCs, previously isolated and cryopreserved, were revived and cultured for seven days in Advanced DMEM (Gibco #12491) supplemented with Glutamax (Invitrogen #35050-061), 10% FBS, and penicillin/streptomycin. Following this period, cells were transferred to a serum-free medium and incubated for an additional three days. The conditioned medium was subsequently harvested, and cells were detached using trypsin and quantified. The conditioned medium was then centrifuged at 500xg for five minutes to remove large particulates. The resulting supernatant was filtered through a 0.2 μm filter to eliminate residual debris. Further purification was achieved using an Amicon® Ultra Centrifugal Filter (30 kDa MWCO), with centrifugation at 4,000xg for 30-60 minutes at 4°C. The isolated extracellular vesicles were then diluted in ratios of 1:10,000 and 1:100,000 and characterized using a ZetaView® particle tracking analysis. Additionally, the extracellular vesicle preparations were evaluated using the Exo-Check™ Exosome Antibody Array according to the manufacturer's instructions.

<u>PBMC Isolation and Culture</u>

Peripheral blood mononuclear cells (PBMCs) were isolated from healthy White Yorkshire x Landrace swine (Oak Hill Genetics, Ewing, IL) that were distinct from the swine used for MSC cultivation. Approximately 8 mL of whole blood was collected and added to Ficoll CPT Vacutainers (BD, Ref 362761), followed by centrifugation at 1,500xg for 28 minutes at 4°C. As per the manufacturer's protocol, the PBMC layer was collected, filtered through a 40 μm filter into a 50 mL Falcon tube, and washed with 20 mL of PBS. The filtered PBMCs were then centrifuged at 500xg for 5 minutes and washed with PBS. Before plating, isolated PBMCs were treated with red blood cell (RBC) lysis buffer for 5 minutes at room temperature, followed by centrifugation and PBS washing. Purified PBMCs were resuspended in RPMI 1640 supplemented with 1x penicillin/streptomycin, 1x sodium pyruvate, and 10% fetal bovine serum (FBS). The cells were cultured at 37°C in a humidified atmosphere of 5% CO2 and ambient oxygen for one week, with bi-daily media changes. After one week, the cells were subjected to activation assays. After aspiration of the culture medium and washing with PBS, naïve PBMCs were activated for 24 hours with either 1× LPS or 1 μg/mL mtDNA with Lipofectamine 2000 Transfection Reagent (TR; Invitrogen, Ref 11668027), as previously described (40) (**Figure S1**) . During this activation period, MSC-EVs labeled with Vybrant DiD Lipophillic Dye (Thermo, Ref V22887) were added in place of FBS. At the end of the 24-hour stimulation period, the conditioned medium was harvested and stored at −20°C for subsequent ELISA and ROS/RNS assays, while the cells were collected for flow cytometric analysis or lysed in Qiazol Reagent for downstream qPCR.

<u>Flow Cytometry</u>

Following a 24-hour activation period, cultured leukocytes were labeled for porcine-specific inflammatory cell-surface markers (41-44) and viability was assessed by 7AAD staining. Flow cytometric assessment was performed on a BD LSRFortessa™ Cell Analyzer and analyzed via De Novo Software FCS Express 7 Plus (version 7.16.0035) following consistent gating strategy implementation (**Figure S2)** to quantify sub-populations of inflammatory granulocytes (CD172+CD163-), inflammatory dendritic cells (CD172+CD16+), and inflammatory macrophages (CD14+CD163+) (41-44) (**Table S1**). Additional staining of MSC-EVs with DiD Lipophillic Dye allowed the detection of MSC-EV uptake by PBMC populations.

Quantitative Polymerase Chain Reaction (qPCR)

RNA was extracted from thawed cell lysate via RNeasy Kit (Qiagen), per the manufacturer's protocol. A NanoDrop spectrophotometer (Thermo Scientific) was used to determine RNA purity and concentration. Total RNA was reverse transcribed and analyzed by qPCR as previously described (45). SsoAdvanced Universal SYBR Green Supermix (BIO-RAD, Ref 1725270) was used in combination with primers for IFNα, IL-1α, IL-1β, IL-6, IL-8, TNFα, and beta-2-microglobulin (β2M) (**Table S2**). The comparative cycle threshold method was used to determine the mRNA expression for each target gene using the gene for β2M as the reference.

Quantification of Conditioned-Media Levels of Inflammatory Cytokines and Reactive Oxygen/Nitrogen Species

Conditioned media (CM) was collected 24-hours post-activation from the supernatant of each cell culture dish well. CM samples were clarified by centrifugation at 500xg for 5-mins to remove cellular debris. CM was thawed and quantified in duplicate with porcine-specific ELISA kits according to manufacturer’s instructions (**Table S3**). Additionally, CM levels of reactive oxygen and nitrogen species (ROS/RNS) were measured using a DCF-based kit (Abcam, Ref ab238535) following the manufacturer's recommendations and as previously described (46, 47).

Statistical Analysis

Data are reported as mean ± standard error of the mean (SEM). Two-tailed unpaired Student’s t-test assessed between-group differences in endpoints measured at single time points. For dose-dependent inhibition curves, simple linear regressions were utilized to determine the 50% inhibitory dose (IC50) and the coefficient of determination (R2). Significance was set to an alpha level of 0.05. All data were compiled with Microsoft Excel (version 2402), while analysis and plotting were performed with GraphPad Prism Software (version 10.2.0).

### SUPPLEMENTAL FIGURES

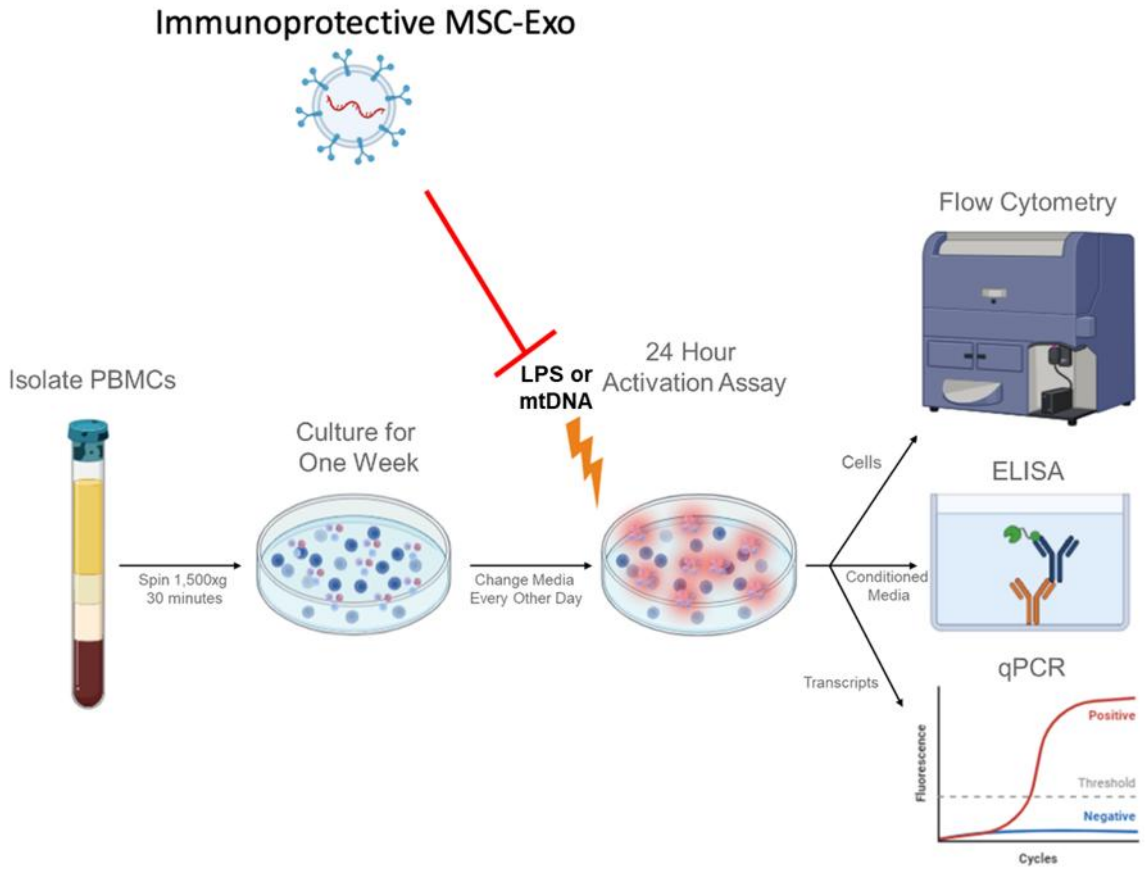


**Figure S1: Experimental Design.** PBMC activation assay experimental design and endpoints following 24hr coculture with MSC-EVs and activation with LPS or mtDNA. Created in BioRender.

### Gating Strategy

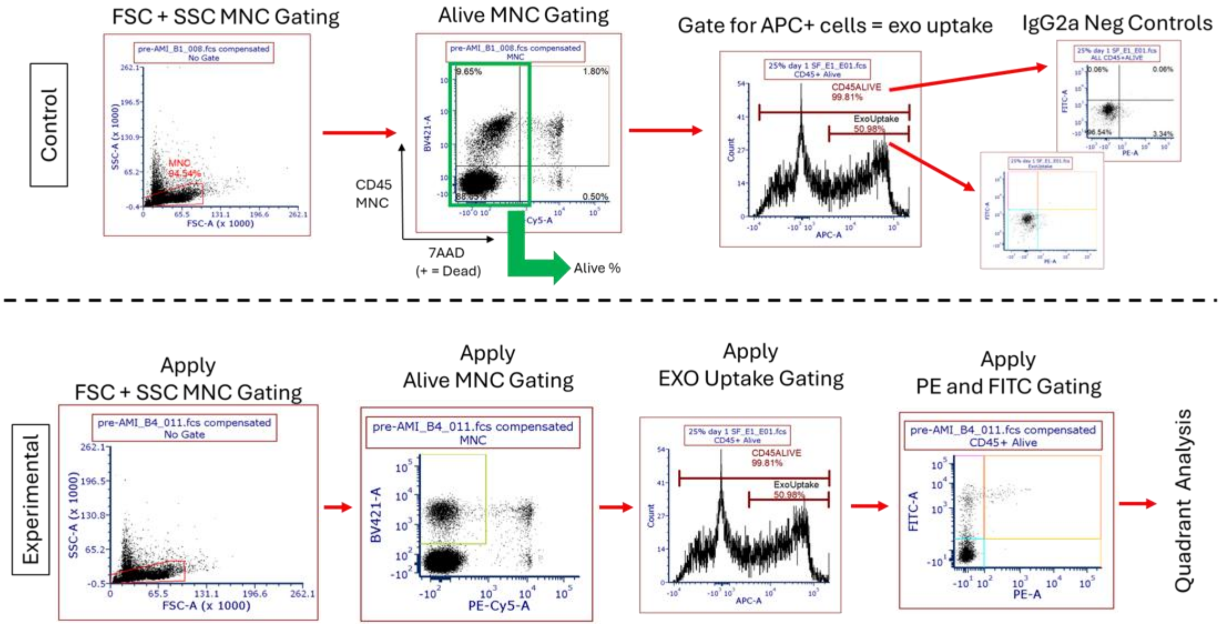


**Figure S2: Flow Cytometry Gating Protocol.** Representative workflow of flow cytometry analysis of PBMCs with uptake of DiD-labeled MSC-EVs.

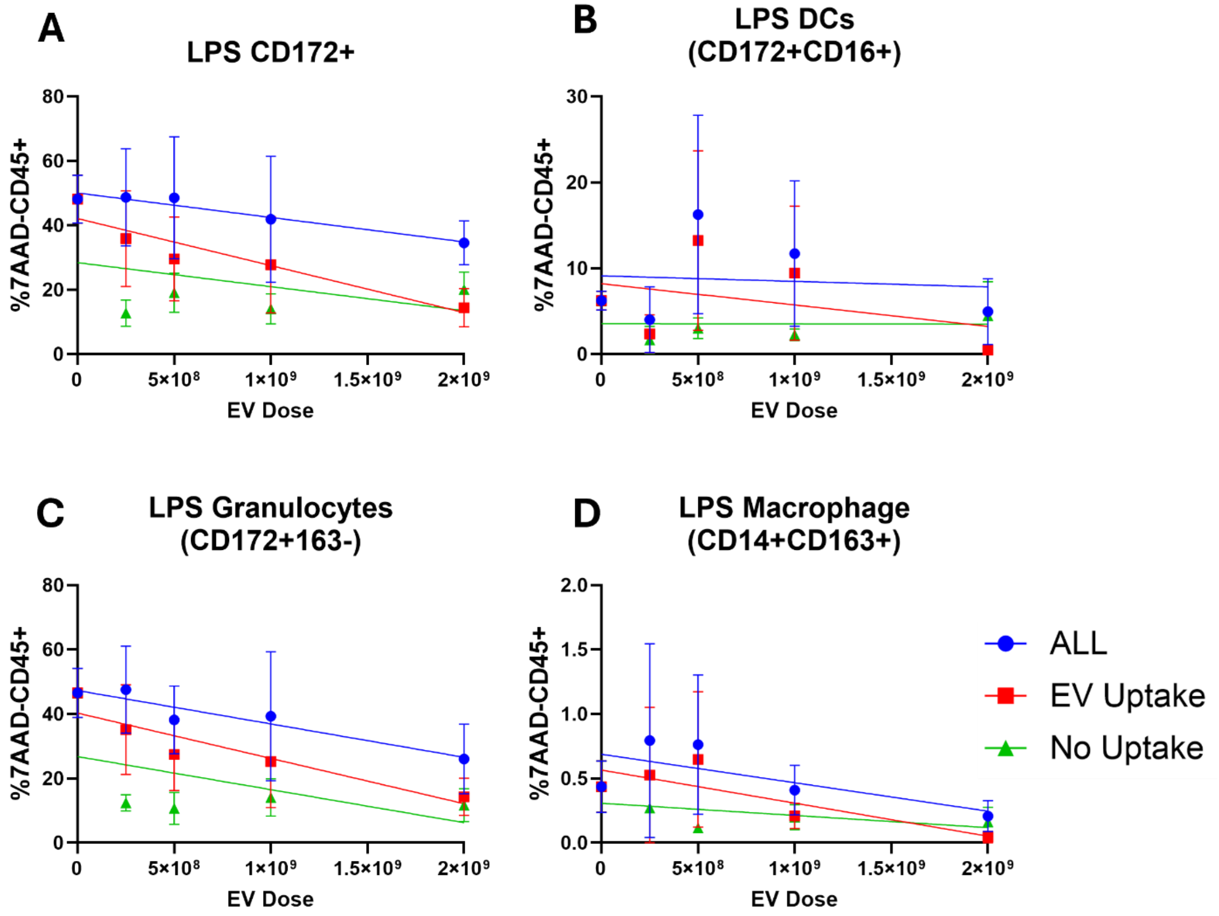


**Figure S3: Dose-Dependent MSC-EVs Attenuation of LPS-Induced Inflammatory Shifts in CD172+ Cells, Dendritic Cells, Granulocytes, and Macrophages.** Relative porcine PBMC (%7AAD-/CD45+) surface marker expression of A) inflammatory adhesion molecule (CD172+) and inflammatory B) dendritic cells (CD172+CD16+), C) granulocytes (CD172+CD163-) and D) macrophages (CD14+CD163+) following co-culture with LPS alone (EV Dose = 0) or LPS treated with either +2.5e8 MSC-EVs, +5e8 MSC-EVs, +1e9 MSC-EVs, or +2e9 MSC-EVs. Data presented as Mean ± SEM. n=biological triplicates, technical duplicates.

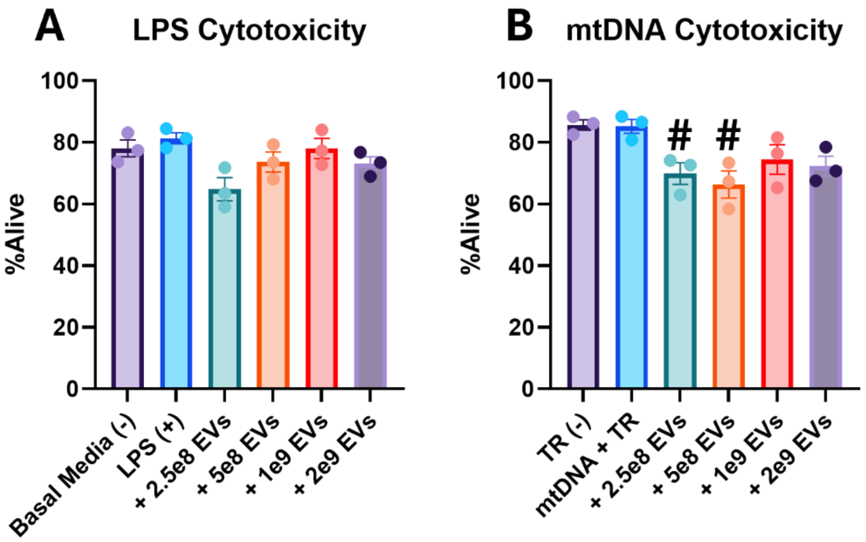


**Figure S4. Cytotoxicity of LPS and mtDNA With and Without MSC-EV Treatment.** Relative % of 7AAD- PBMCs after 24hr coculture with A) LPS or B) mtDNA+TR and either 2.5e8 MSC-EVs, 5e8 MSC-EVs, 1e9 MSC-EVs, or +2e9 MSC-EVs. Mean ± SEM. n=biological triplicates. #$p<0.05$ vs (-) control.

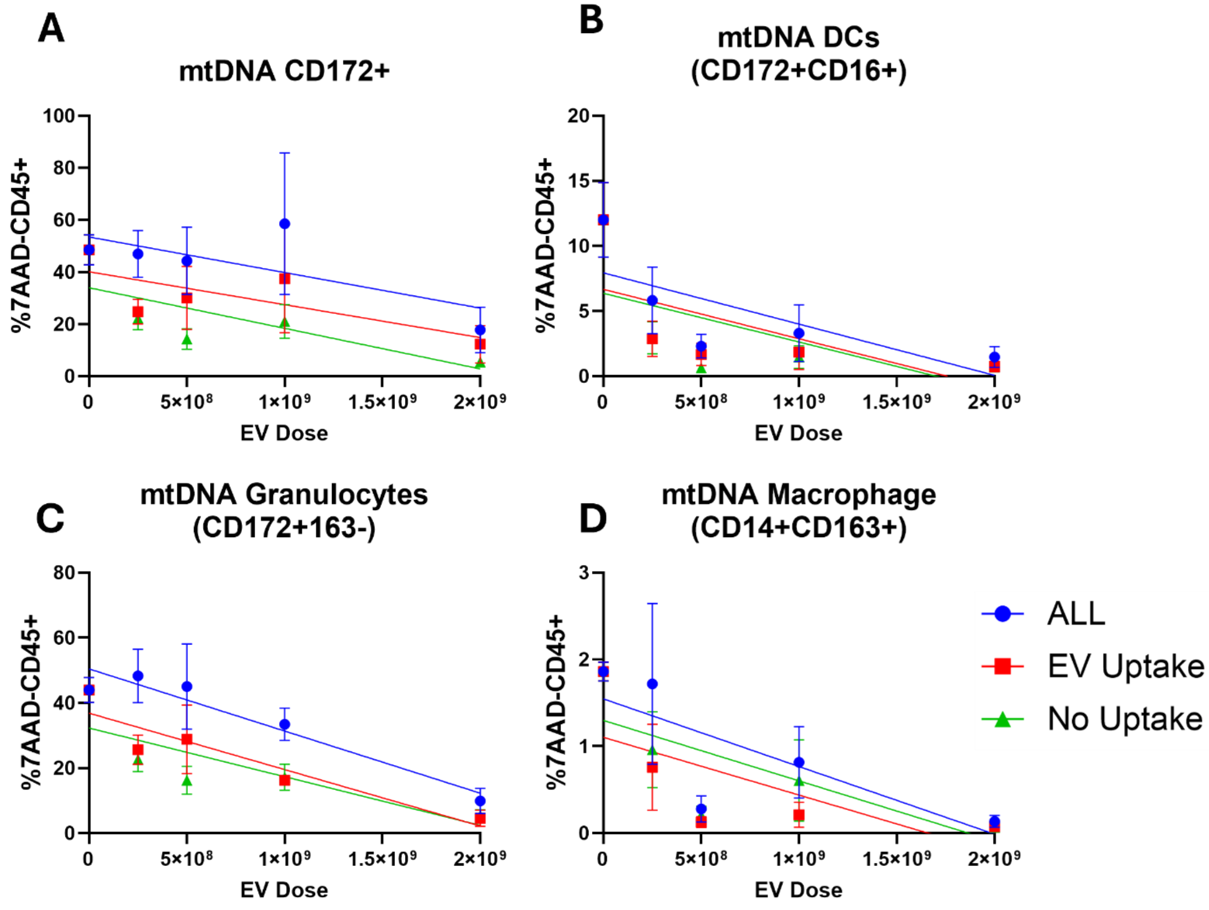


**Figure S5: Dose-Dependent MSC-EVs Attenuation of mtDNA-Induced Inflammatory Shifts in CD172+ Cells, Dendritic Cells, Granulocytes, and Macrophages.** Relative porcine PBMC (%7AAD-/CD45+) surface marker expression of A) inflammatory adhesion molecule (CD172+) and inflammatory B) dendritic cells (CD172+CD16+), C) granulocytes (CD172+CD163-) and D) macrophages (CD14+CD163+) following co-culture with mtDNA+TR alone (EV Dose = 0) or mtDNA+TR treated with either +2.5e8 MSC-EVs, +5e8 MSC-EVs, +1e9 MSC-EVs, or +2e9 MSC-EVs. Data presented as Mean ± SEM. n=biological triplicates, technical duplicates.

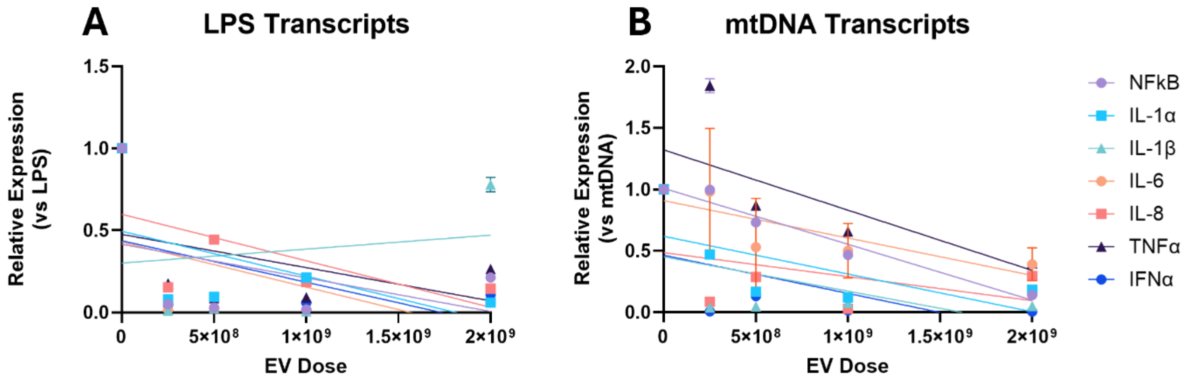


**Figure S6: Dose-Dependent Response of MSC-EVs Against LPS and mtDNA Induction of Inflammatory Gene Expression.** Relative PBMC inflammatory transcript expression of IFNα, IL-1α, IL-1β, IL-6, IL-8, and TNFα at 24-hr post-activation assay with A) LPS or B) mtDNA+TR and treated with either +2.5e8 MSC-EVs, +5e8 MSC-EVs, +1e9 MSC-EVs, or +2e9 MSC-EVs. Housekeeping gene = β2M. Data presented as Mean ± SEM. n=biological triplicates, technical duplicates.

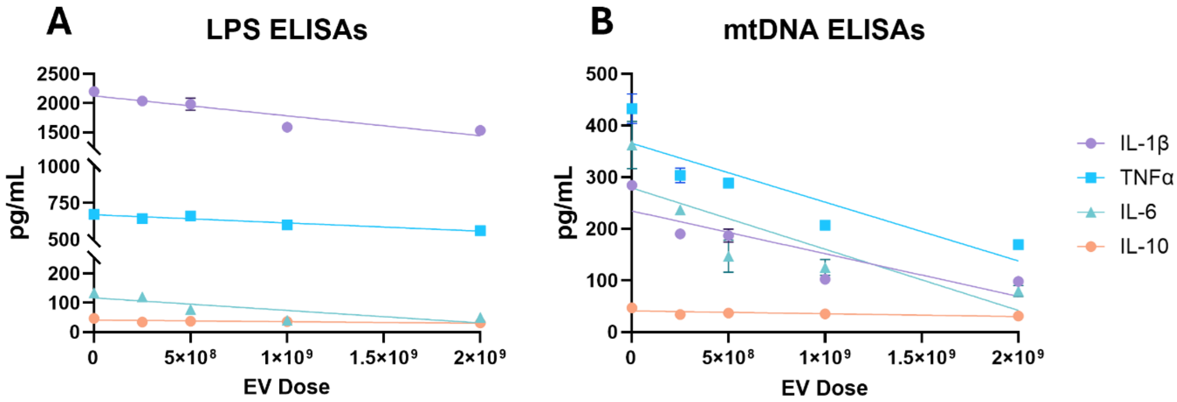


**Figure S7: Dose-Dependent Response of MSC-EVs Against Inflammatory Cytokine Release From LPS- and mtDNA-Activated PBMCs.** Conditioned media levels of PBMC-secreted inflammatory cytokines TNFα, IL-1β, IL-6, and IL-10 following a 24hr activation assay with A) LPS or B) mtDNA+TR and treated with +2.5e8 MSC-EVs, +5e8 MSC-EVs, +1e9 MSC-EVs, or +2e9 MSC-EVs. Data presented as Mean ± SEM. n=biological triplicates.

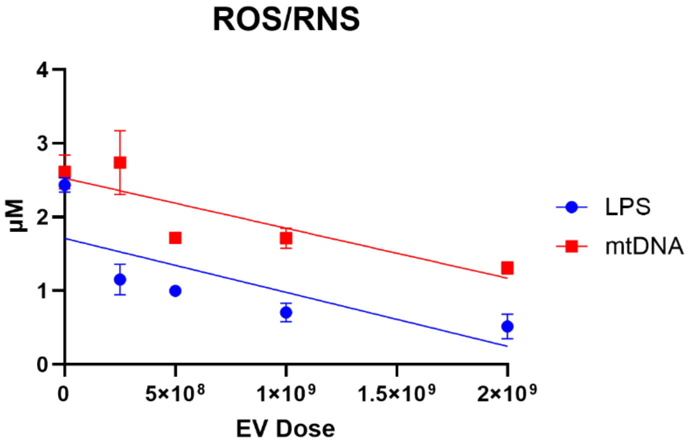


**Figure S8: Dose-Dependent Response of MSC-EVs to Attenuate LPS- and mtDNA-Induced ROS and RNS Production.** Conditioned media levels of reactive oxygen and nitrogen species (ROS/RNS) following a 24 hr post-activation assay with A) LPS or B) mtDNA+TR and treated with either +2.5e8 MSC-EVs, +5e8 MSC-EVs, +1e9 MSC-EVs, or +2e9 MSC-EVs against transfection reagent (TR; negative control). Data presented as Mean ± SEM. n=biological triplicates, technical duplicates. *p<0.05 vs. mtDNA+TR. #p<0.05 vs. Basal Media (-) or TR (-).

### SUPPLEMENTAL TABLES

| Antibody | Vender | Catalog # | Dilution |
|---|---|---|---|
| Fc Block | Miltenyi Biotec | 130-059-901 | 2.5ug/1e6 cells |
| 7AAD | Invitrogen | 00-6993-50 | 0.25ug/1e6 cells |
| αIgG1 (FITC) | Bio-rad | MCA928F | 0.25uL/1e6 cells |
| αIgG1 (PE) | Bio-rad | MCA928PE | 0.25uL/1e6 cells |
| αIgG2a (PE) | R&D Systems | IC003P | 0.25uL/1e6 cells |
| αIgG2b (FITC) | Bio-rad | MCA691F | 0.25uL/1e6 cells |
| αCD45 (PB) | Bio-rad | MCA1222PB | 0.25uL/1e6 cells |
| αCD172 (FITC) | BD Biosciences | 561498 | 0.5ug/1e6 cells |
| αCD163 (PE) | Bio-rad | MCA2311PE | 0.25uL/1e6 cells |
| αCD14 (FITC) | Bio-rad | MCA1218F | 0.25uL/1e6 cells |
| αCD16 (PE) | Bio-rad | MCA1971PE | 0.25uL/1e6 cells |

**Table S1. Flow Cytometry Antibodies for Surface-Antigen Detection of Porcine Innate Immune Cells.**

| Gene | Accession | FWD (5' ->3') | REV (5'-3') |
|---|---|---|---|
| IFNα | AC127471 | GGCTCTGGTGCATGAGATGT | GCCTTCTTCCTGAATCTGTCTTA |
| IL-1α | P18430 | TATGCCTCTGAGTACCTCTAA | TCGTCATCGGTGATGAACTAA |
| IL-1β | NC_010445.4 | TGCTATCATCTCCTTGCACA | ACGATAGTAGAGGAACGTGT |
| IL-6 | NC_010451.4 | CCTGAGATTGATGCCGTCCA | TCACAGGATTGCGAGTATGAAA |
| IL-8 | X61151.1 | GCATAAATACGCATTCCACA | CAACAACAACGAAGAGTCAAGAG |
| TNFα | NC_010449.5 | AGAGTGGGTATGCCAATGCC | ACTACCACACTCACTCCTTTTG |
| β2M | NM_213978 | AAACGGAAAGCCAAATTACC | ATCCACAGCGTTAGGAGTGA |

**Table S2. Oligonucleotide primer sequences for qPCR detection of porcine genes.**

| ELISA Kit | Vender | Catalog # | Target (Species) |
|---|---|---|---|
| TNFα | R&D Systems | PTA00 | Porcine |
| IL-1β/IL-1F2 | R&D Systems | PLB00B | Porcine |
| IL-6 | R&D Systems | P6000B | Porcine |
| IL-10 | R&D Systems | P1000 | Porcine |

**Table S3. ELISA Kits of Porcine Cytokine Quantification.**

| FLOW | | LPS | | | mtDNA | | |
|---|---|---|---|---|---|---|---|
| | | ALL | EV Uptake | No Uptake | ALL | EV Uptake | No Uptake |
| Inf. Adhesion | IC50 | 5.22E+09 | 4.25E+09 | 2.74E+10 | 2.14E+09 | 3.30E+09 | 1.39E+09 |
| | R2 | 0.002 | 0.270 | 0.529 | 0.504 | 0.210 | 0.733 |
| | p-value | 0.382 | 0.050* | 0.163 | 0.153 | 0.101 | 0.001* |
| Inf. Dendritic Cells | IC50 | 1.05E+10 | 3.61E+09 | --- | 5.54E+08 | 6.19E+08 | 4.88E+08 |
| | R2 | 0.008 | 0.047 | 0.448 | 0.539 | 0.586 | 0.491 |
| | p-value | 0.881 | 0.512 | 0.980 | 0.015* | 0.014* | 0.017* |
| Inf. Granulocytes | IC50 | 3.68E+09 | 3.81E+09 | 3.30E+09 | 1.49E+09 | 1.54E+09 | 1.43E+09 |
| | R2 | 0.903 | 0.388 | 0.098 | 0.923 | 0.786 | 0.906 |
| | p-value | 0.193 | 0.045* | 0.055 | <0.001* | <0.001* | <0.001* |
| Inf. Macrophages | IC50 | 2.23E+09 | 1.95E+09 | 3.55E+09 | 8.20E+08 | 6.63E+08 | 9.68E+08 |
| | R2 | 0.471 | 0.355 | 0.160 | 0.615 | 0.594 | 0.597 |
| | p-value | 0.402 | 0.226 | 0.320 | 0.021* | 0.006* | <0.006* |

**Table S4. Flow Cytometry of Inflammatory Leukocyte Sub-Populations Following Increasing MSC-EV Concentrations.**

**Abbreviations:** *, significant relationship; EV, extracellular vesicle; IC50, inhibitory concentration of 50%; R2, R-squared; Inf., inflammatory.

| qPCR | | LPS | mtDNA |
|---|---|---|---|
| NFκB | IC50 | 1.02E+09 | 1.11E+09 |
| | R2 | 0.149 | 0.959 |
| | p-value | 0.156 | <0.001* |
| IL-1α | IC50 | 9.05E+08 | 1.01E+09 |
| | R2 | 0.291 | 0.433 |
| | p-value | 0.038* | <0.001* |
| IL-1β | IC50 | --- | 8.10E+08 |
| | R2 | 0.019 | 0.271 |
| | p-value | 0.622 | 0.046* |
| IL-6 | IC50 | 7.82E+08 | 1.73E+09 |
| | R2 | 0.249 | 0.222 |
| | p-value | 0.058 | 0.119 |
| IL-8 | IC50 | 1.06E+09 | 1.25E+09 |
| | R2 | 0.377 | 0.158 |
| | | 0.015* | 0.142 |
| TNFα | IC50 | 1.18E+09 | 1.35E+09 |
| | R2 | 0.173 | 0.504 |
| | p-value | 0.123 | 0.003* |
| IFNα | IC50 | 8.67E+08 | 7.49E+08 |
| | R2 | 0.223 | 0.320 |
| | p-value | 0.076 | 0.028* |

**Table S5. Inflammatory Transcript Following Increasing MSC-EV Concentrations.**

**Abbreviations:** *, significant relationship; IC50, inhibitory concentration of 50%; R2, R-squared; Inf., inflammatory.

| ELISA + ROS/RNS | | LPS | mtDNA |
|---|---|---|---|
| TNFα | IC50 | 5.97E+09 | 1.61E+09 |
| | R2 | 0.920 | 0.782 |
| | p-value | <0.001* | <0.001* |
| IL-6 | IC50 | 1.36E+09 | 1.18E+09 |
| | R2 | 0.661 | 0.700 |
| | p-value | <0.001* | <0.001* |
| IL-1β | IC50 | 3.12E+09 | 1.42E+09 |
| | R2 | 0.846 | 0.728 |
| | p-value | <0.001* | <0.001* |
| IL-10 | IC50 | 3.38E+09 | 3.76E+09 |
| | R2 | 0.741 | 0.530 |
| | p-value | 0.018* | 0.017* |
| ROS/RNS | IC50 | 1.17E+09 | 1.86E+09 |
| | R2 | 0.589 | 0.736 |
| | p-value | 0.002* | 0.001* |

**Table S6. Cytokines and ROS/RNS in Conditioned Media Following Increasing MSC-EV Concentrations.**

**Abbreviations:** *, significant relationship; EV, extracellular vesicle; IC50, inhibitory concentration of 50%; R2, R-squared; Inf., inflammatory.
